# Development of novel anti-malarial from structurally diverse library of molecules, targeting plant-like Calcium Dependent Protein Kinase 1, a multistage growth regulator of *P. falciparum*

**DOI:** 10.1101/2020.01.14.907147

**Authors:** Ravi Jain, Sakshi Gupta, Manoj Munde, Soumya Pati, Shailja Singh

## Abstract

Upon *Plasmodium falciparum* merozoites exposure to low [K^+^] environment in blood plasma, there is escalation of cytosolic [Ca^2+^] which activates Ca^2+^-Dependent Protein Kinase 1 (CDPK1), a signaling hub of intra-erythrocytic proliferative stages of parasite. Given its high abundance and multidimensional attributes in parasite life-cycle, this is a lucrative target for desiging antimalarials. Towards this, we have virtually screened MyriaScreenII diversity collection of 10,000 drug-like molecules, which resulted in 18 compounds complementing ATP-binding pocket of CDPK1. *In vitro* screening for toxicity in mammalian cells revealed that these compounds are non-toxic in nature. Further, SPR analysis demonstrated differential binding affinity of these compounds towards recombinantly purified CDPK1 protein. Selection of lead compound 1 was performed by evaluating their inhibitory effects on phosphorylation and ATP binding activities of CDPK1. Further, *in vitro* biophysical evaluations by ITC and FS revealed that binding of compound 1 is driven by formation of energetically favorable non-covalent interactions, with different binding constants in presence and absence of Ca^2+^, and TSA authenticated stability of compound 1 bound CDPK1 complex. Finally, compound 1 strongly inhibited intra-erythrocytic growth of *P. falciparum in vitro*. Concievably, we propose a novel CDPK1-selective inhibitor, step towards developing pan-CDPK kinase inhibitors, prerequisite for cross-stage anti-malarial protection.

## Introduction

*Plasmodium falciparum* is the most prevalent malaria parasite which accounted for majority of the total estimated malaria cases in WHO African Region (99.7%), the Western Pacific (71.9%), the Eastern Mediterranean (69%) and WHO region of Southeast Asia (62.8%) (statistics are as per 2017). (1) Five countries in WHO sub-Saharan African Region and *Republic of India* have been reported to carry almost half of the global malaria burden. Children aged under 5 years constitute the most vulnerable group of the population which accounted for almost 61% of all malaria deaths worldwide in 2017. (1) Clinical manifestations of malaria largely result from exponential expansion of the parasite in blood wherein parasites invade into, multiply within and evade from erythrocytes for their propagation. Currently, there is no drug regime available that can target all the intra-erythrocytic proliferation stages of the parasite.

In this process of parasitic growth, exposure of *P. falciparum* merozoites from high (host cytosolic; 140 mM) to low (blood plasmic; 5.4 mM) potassium-ion (K^+^) environment triggers phospholipase-C mediated intracellular Ca^2+^ release from endoplasmic reticulum. (2) This escalation of cytosolic Ca^2+^ activates Calcium-Dependent Protein Kinases (CDPKs) of the parasite. In the genus Plasmodium, CDPK proteins are present as a multigene family comprised of seven members and each gene is predominantly expressed in a distinct phase of the parasite life cycle. For example, CDPK3 is expressed in ookinetes and *P. berghei* parasites without a functional *cdpk3* gene show reduced ability to invade midgut of the mosquito host, while simultaneously showing reduction in the gliding motility phenotype. (3, 4) Similarly, *Pb*CDPK4 regulates sexual differentiation into male gametocytes and therefore, being crucial for transmission to the mosquito host and sexual reproduction. (5) Likewise, *Pf*CDPK5 guides egress of merozoites from the host erythrocytes. (6) However, besides these members of *cdpk* multigene family (7, 8), one of the most widely studied member is CDPK1, which is a signaling hub responsible for multistage cellular processes involved in schizogony, merozoite invasion, gametogenesis, and transmission to the mosquito host. (9–13)

Mechanistic studies revealed that Protein Kinase G (PKG)-mediated phosphorylation of CDPK1 allows its localization to apical structures. (14) Another piece of study also depicted that CDPK1 regulates actin-myosin motor during gliding and invasion of the parasite by phosphorylating a 50-kDa Glideosome-Associated Protein (GAP-45) and Myosin Tail domain Interacting Protein (MTIP), two important components of Inner Membrane Complex (IMC) of *P falciparum* merozoites. Along with the proteins of IMC, it also phosphorylates regulatory subunit of cAMP-dependent Protein Kinase A (PKAr), thereby regulating cAMP-mediated signaling cascade module. (13) With this understanding of an indispensable role of CDPK1 in the parasite life cycle, we pursued this as a potential target for development of novel *specific* inhibitors that can block parasite growth targeting multiple stages in malarial infection. In this study, we have identified small molecule ligands that bind to and inhibit the functional (kinase) activity of CDPK1 *in vitro* and parasite growth *ex vivo*, by combining high-throughput virtual screening strategy, followed by experimental evaluation by adopting several biochemical and biophysical assays *in vitro*. This study can in principle be extended towards developing novel drug candidates against any eukaryotic parasite by systematic high-throughput screening of small molecule-based drug-like libraries.

## Materials and Methods

### Culture of *P. falciparum* 3D7 and HepG2

*P. falciparum* strain 3D7 was cultured according to the previously described protocol. (15) Briefly, parasite culture was maintained in O+ erythrocytes at 4% hematocrit in RPMI 1640 medium (Gibco®, USA), supplemented with 0.5% AlbuMAX^TM^ I (Gibco®, USA), 50 mg/L hypoxanthine, 10 mg/L gentamycin (Gibco®, USA) and 2 gm/L sodium bicarbonate. Culture was maintained in ambient hypoxic environment (5% O_2_, 5% CO_2_, balanced with nitrogen).

The human hepatocellular carcinoma cell line, HepG2 was cultured in Dulbecco’s Modified Eagle’s Medium (Gibco®, USA) supplemented with 10% fetal bovine serum (Gibco®, USA) and penicillin-streptomycin (100 units/ml and 100 mg/ml, respectively) solution. Culture was maintained at 37°C in a humidified atmosphere containing 5% CO_2_.

### Purification of 6xHis-CDPK1, Homology modeling and Virtual screening

Cloning, over-expression and purification of 6xHis-CDPK1 was done as described previously. (11) To raise anti-sera, mice were immunized by injecting the emulsified protein. Comparative or homology modeling was employed to model 3D structure of CDPK1 from *P. falciparum* strain 3D7. To accomplish this, amino acid sequence of PfCDPK1 protein (PF3D7_0217500) was retrieved from PlasmoDB database. (16) To search for a suitable template for homology modeling, BLASTp search was performed using amino acid sequence of PfCDPK1 as query sequence, against Protein Data Bank (PDB) database. (17, 18) Amino acid sequence identity between CDPK1 orthologs from *P. falciparum* 3D7 and *P. berghei* strain ANKA was found to be 93%, thus rendering X-Ray diffraction based structural model of *Pb*CDPK1 (PDB ID: 3Q5I; resolution: 2.1 Å) as a suitable template to model 3D structure of PfCDPK1. (19) Homology modeling was done by using Modeller v9.17 software. (20) Generated model was further subjected to structural refinement by using ModRefiner. (21) Reliability of the refined structural model of CDPK1 was assessed by examining backbone dihedral angles: phi (Ø) and psi (ψ) of the amino acid residues lying in the energetically favourable regions of Ramachandran space. (22) This was done by using PROCHECK v.3.5. (23) Percentage quality assessment of the protein structure was evaluated by utilizing four sorts of occupancies: core, additional allowed, generously allowed and disallowed regions. The refined 3D structural model of PfCDPK1, thus generated, was subsequently used for docking studies.

Novel small molecule inhibitors of CDPK1 were identified by virtual screening of MyriaScreen II diversity collection of 10,000 drug-like screening compounds. (24) Structural Data Format (SDF) file of Myriascreen II library was requested from Sigma-Aldrich. Ligands were subjected to virtual screening against 3D structural model of CDPK1 using Autodock Vina Tools v1.5.6. (25) This was done by constructing a virtual 3D grid covering the entire protein structure, with default spacing and high exhaustiveness score. Top ranked compounds were selected based on the highest (most negative) binding affinity to the catalytically active site (ATP binding pocket) of CDPK1. Finally, compounds binding to the ATP binding pocket of CDPK1 were further filtered on the basis of their propensity to follow Lipinski’s rule by using Drug Likeness Tool (DruLiTo). (26) The PyMOL Molecular Graphics System, v2.1 by Schrödinger, LLC was used for molecular visualization of CDPK1 structural model and AutoDock Vina results. (27)

### Evaluation of cytotoxicity in HepG2 cell line

Cytotoxic effect on HepG2 was assessed by utilizing ability of live cells to cleave MTT [(3-[4,5-dimethylthiazol-2-yl]-2,5-diphenyl tetrazolium bromide)] into blue colored formazan crystals. Briefly, HepG2 cells were seeded in a 96-well microtiter plate at a density of 10,000 cells per 100 μL per well. Cells were allowed to proliferate at 37°C for 48 hrs in the presence of compounds at 100 μM concentration. Cells cultured in the absence of any compound were maintained as positive control. After treatment, MTT labeling reagent was added to each well to a final concentration of 0.5 mg/ml and incubated at 37°C for 4 hrs. Purple colored formazan crystals, thus formed, were dissolved in 100 ul DMSO solvent and optical density was measured spectrophotometrically by taking absorbance at 570 nm using Varioskan™ LUX multimode microplate reader (Thermo Fisher Scientific™). The absorbance of untreated cells was taken as 100%.

### Surface Plasmon Resonance (SPR) analysis of binding affinity between compounds and CDPK1

To determine binding strength of compounds (N=18) procured from virtual screening of MyriaScreen II library, real-time biomolecular interaction analysis with SPR was carried out at physiologically relevant concentrations by using AutoLab Esprit SPR. Kinetic rate constants: K_a_ and K_d_ (association and dissociation rate constants, respectively) as well as affinity constant, K_D_ were measured at room temperature (RT, 293K). SPR analysis was performed by following previously described protocols. (28) Briefly, interaction kinetics was studied by injecting compounds at different concentrations: 100, 75, 50, 25 and 12.5 μM over the CDPK1 (7.43 ng per mm^2^) immobilized SPR sensor chip surface, followed by comparing their respective kinetics & binding affinities at RT. A control flow cell was activated and blocked in a similar manner with injection of equivalent concentrations of DMSO to allow for reference subtraction. HEPES-NaOH buffer was used both as immobilization and binding solutions. Data were fit to the two-state conformational change model using AutoLab SPR Kinetic Evaluation software provided with the instrument. K_D_ value was calculated using the Integrated Rate Law (IRL) equation. Two independent experiments were performed.

### *In vitro* kinase assay with CDPK1

Functional activity of 6xHis-CDPK1 was monitored by utilizing ADP-Glo™ Kinase Assay (Promega Corporation), which is ATP regeneration based luciferase reaction system resulting from nascent ADP phosphorylation. The luminescence signal, thus generated, is proportional to the amount of ADP released in a given kinase reaction. Phosphorylation experiments were performed with 100 ng of 6xHis-CDPK1, per reaction, in assay buffer (100 mM Tris-Cl, pH 7.4; 2.5 mM DTT; 50 mM MgCl_2_ and 2.5 mM MnCl_2_), by following previously described protocol. (29) Enzymatic reaction was carried out in the absence and presence of calcium ions. 10 μg of dephosphorylated casein from bovine milk, per reaction, was used as exogenous substrate for the enzyme. 2.5 mM EGTA was added for conditions requiring absence of Ca^2+^ ions. Kinase reactions were initiated by adding 1 μM ATP and allowed to take place at 30°C for 1 hr. To test for any inhibitory effect of compounds (N=18) screened from MyriaScreen II Diversity Collection, CDPK1 was allowed to pre-incubate in the reaction buffer with 50 μM of each compound at room temperature for 1 hr, followed by initiating the reactions by adding ATP. Upon completion of the reactions, ADP-Glo™ Kinase Assay was performed. Luminescence signal, thus generated, was measured with Lumat³ LB 9508 Ultra-Sensitive Tube Luminometer (Berthold Technologies, USA). Kinase activity of CDPK1 was further assessed in the presence of different concentrations of lead compounds: ST092793 (assigned as compound 1) and S344699 (assigned as compound 2) to evaluate their respective IC_50_ values. All experiments were done at least thrice.

Phosphorylation status of CDPK1 and casein in presence of the lead compounds 1_(ST092793)_ and 2_(S344699)_ was checked by *in-gel* fluorescence staining with phosphoamino acid stain, Pro-Q Diamond (Thermo Fisher Scientific™). Protocol was followed as recommended by the supplier. Briefly, SDS-PAGE was performed for the phosphorylation reactions carried out in the presence of different concentrations: 100, 50, 25 and 12.5 μM of lead compounds 1_(ST092793)_ and 2_(S344699)_. Phosphostained gels were imaged using Typhoon™ FLA 9500 biomolecular imager (GE Healthcare Life Sciences). Excitation was set at 555 nm with emission filter of 580 BP 30. After visualization, gel(s) was stained with Coomassie Brilliant blue G-250 dye to confirm equal loading of CDPK1 and dephosphorylated casein in all the lanes. Dried coomassie gels were scanned on Epson CX5400 high resolution scanner. Estimates of band intensities from both Pro-Q Diamond and Coomassie stained gel images were made using ImageJ 1.52a. The experiment was done thrice.

### Competitive ATP affinity chromatography assay

To confirm if the lead compounds 1_(ST092793)_ and 2_(S344699)_ inhibit functional activity of CDPK1 by directly binding to its active site (ATP-binding pocket), affinity chromatography of CDPK1 pre-incubated with the compounds was performed by utilizing ATP Affinity Test Kit (Jena Bioscience). Protocol was followed as recommended by the manufacturer. Briefly, 10 μg of purified CDPK1 protein per reaction was incubated with 100 μM of compound 1_(ST092793)_ and/or 2_(S344699)_ in binding solution for 1 hr at RT. This interaction was competed with 8AH-ATP-agarose {8-[(6-Amino) Hexyl]-amino-ATP-agarose} in which ATP is immobilized on the agarose matrix via the adenine base through C8 linker. Blank agarose served as negative control. Elution fractions were analyzed by SDS-PAGE and immunobloting by probing recombinant CDPK1 with monoclonal poly-Histidine-peroxidase antibody. CDPK1 present in flowthroughs of control and test samples served as loading controls for normalization. Signals were detected with SuperSignal™ West Femto Maximum Sensitivity Substrate (Thermo Fisher Scientific™). Estimates of band intensities of CDPK1 from both control and treated reactions were made using ImageJ 1.52a. The experiment was done twice.

### Isothermal Titration Calorimetry (ITC) and Steady-state fluorescence spectroscopy

To calculate kinetic parameters such as binding affinity constant, K_a_ for the interaction of CDPK1 with *relatively more potent inhibitor* (i.e., compound 1_(ST092793)_), ITC experiments were performed by using MicroCal iTC200 (Malvern Instruments Ltd, UK). For this purpose, recombinant CDPK1 protein was dialyzed extensively against HEPES-NaCl buffer (10 mM HEPES-NaOH, pH 7.4 and 150 mM NaCl) using Amicon^TM^ Ultra-15 Centrifugal Filter Unit (10 kDa cutoff) before its subsequent use in ITC. Dilutions of CDPK1 and compound 1_(ST092793)_ were prepared in the same buffer to ignore contribution from buffer‐buffer interaction, and degassed by vacuum for 10 minutes prior to use. ITC analysis was done in the absence and presence of 2.5 mM CaCl_2_ at RT, by following previously described protocols. (30) Briefly, syringe was loaded with 100 μM of the ligand (i.e., compound 1_(ST092793)_), and sample cell was filled with 10 μM of CDPK1. Background titration profiles were obtained by injecting ligand into the buffer under identical experimental conditions. The experiment was done twice. Amount of heat produced per injection (corrected for heat of dilution of the ligand) was analyzed by integration of area under individual peaks by MicroCal ORIGIN 7 software provided by the instrument manufacturer. Experimental data was presented as the corrected amount of heat produced per second (μcal/sec) following each injection of the ligand into the protein solution as a function of time (minutes). The non-linear experimental data was fit to the single-site binding model yielding molar binding stoichiometry (N), binding constant (Ka), enthalpy change (ΔH), entropy change (ΔS) and Gibbs free energy (ΔG).

To confirm the results obtained with ITC, fluorescence spectroscopic analysis was performed by using Cary Eclipse Fluorescence Spectrophotometer (Agilent Technologies, US). All experiments were done at RT using a quartz cell with 1 cm path length. Dilution of CDPK1 was prepared in HEPES-NaCl buffer, in the absence and presence of calcium ions. Intrinsic fluorescence emission spectra were recorded for free (unbound) CDPK1 protein (5 μM) as well as upon its complexation with increasing concentrations of compound 1_(ST092793)_ (0.5 μM to 29 μM, in the absence of calcium; and, 1 μM to 38 μM in the presence of calcium). The emission spectra for CDPK1 were recorded in the range of 300–450 nm after excitation at 295 nm to selectively excite tryptophan residues of the protein. Excitation and emission slits were set at 10 nm. All spectra were corrected by subtracting the corresponding buffer baseline, obtained under similar conditions. The experiment was done twice. Data obtained from CDPK1 quenching experiments were fitted using Kaleidagraph 4.0.

### Thermal Stability Assay (TSA)

Thermostability of CDPK1 was monitored for the detection of binding events between CDPK1 and Compound 1_(ST092793)_ by following the protocol as described earlier. (31) Briefly, 10 μg of the purified recombinant protein per reaction (in HEPES-NaCl buffer, pH 7.5; 30 μL) was incubated with 100 μM of the compound, in the absence and presence of 2.5 mM CaCl_2_. Protein in control reactions was incubated with an equivalent volume of DMSO. Following incubation, reaction mixtures were heated on Mastercycler™ Nexus Thermal Cycler (Eppendorf™) at different temperatures ranging from 40°C to 90°C for 6 min and then cooled down to room temperature for 4 min. Following centrifugation at 16,800g for 40 min at 4°C, SDS-PAGE of supernatants was performed. Estimates of band intensities of CDPK1 protein from Coomassie stained gel images were made using ImageJ 1.52a. For analysis of melting shift, CDPK1 signal intensity was normalized to the respective intensity at 40°C and quantified protein levels were analyzed.

### Evaluation of growth inhibition of malaria parasite

Lead compound 1_(ST092793))_ with potent CDPK1-inhibitory activity was subjected to dose-dependent cytotoxic evaluation on parasite growth at concentrations ranging from 1.56 to 100μM. Briefly, trophozoites at 1% initial parasitaemia were treated with the compounds for one complete intra-erythrocytic life cycle of the parasite. Untreated parasites served as control. Post incubation, parasites were stained with Ethidium Bromide (10 μM), followed by flow cytometry on BD LSRFortessa™ cell analyzer using FlowJo v10 software. Fluorescence signal (FL-2) was detected with the 590 nm band pass filter by using an excitation laser of 488 nm collecting 100,000 cells per sample. Following acquisition, parasitaemia levels were estimated by determining the proportion of FL-2 positive cells using Cell Quest. Percent parasite growth inhibition was calculated as follows: % Parasite growth inhibition = [1 − (% Parasitaemia _(treatment)_ / % Parasitaemia _(Control)_] X 100. IC_50_ value was calculated by using Origin® 2018b Graphing and Analysis Software (OriginLab Corporation).

### Statistical analysis

In the bar graphs, data are expressed as the mean ± standard deviation (s.d.) of three independent experiments done in duplicates. Unless stated, data analysis of the assay plots and calculation of biochemical parameters were done by using both Microsoft Excel and Origin® 2018b Graphing and Analysis Software (OriginLab Corporation) were used. Unless indicated, the differences were considered to be statistically significant at P<0.05.

## Results

### Purification of 6xHis-CDPK1, three-dimensional structural model of *Pf*CDPK1 and its quality assessment

Coomassie-stained gel shows the level of purity of recombinant *Pf*CDPK1 used for all assays (Fig. 1a). Anti-sera raised in mice was used to probe recombinantly purified CDPK1 protein and native protein from schizonts lysate (Fig. 1a). *Pf*CDPK1 contains an N-terminal KD tethered to a C-terminal CamLD via JD (Fig. 1b). Three-dimensional structure of a protein can provide us with precise information about its single, most stable conformation, as dictated by its sequence. X-Ray diffraction based structural model of *Pb*CDPK1 was used as a template to generate 3D coordinates of *Pf*CDPK1 by homology modeling, one of the most common structure prediction methods in structural genomics and proteomics (Fig. S1a). After optimal rigid-body superimposition of *Pb*CDPK1 with the generated structural model of *Pf*CDPK1, overall Root-Mean-Square Deviation (RMSD) value of the C-alpha atomic coordinates was found to be 0.31 Å, suggesting a reliable 3D structure of *Pf*CDPK1 (Fig. S1b). Assessment of stereochemical quality and accuracy of the generated homology model displayed 89.2% of amino acid residues lying in the most favored (“core”) regions, with 8.1%, 1.9%, and 0.8% residues in “additional allowed”, “generously allowed” and “disallowed regions” of Ramachandran plot, respectively (Fig. S1c). Also, protein structure with ≥90% of its amino acid residues lying in the most favored regions of Ramachandran plot is considered to be as accurate as a crystal structure at 2Å-resolution. This indicated that the backbone dihedral angles: phi and psi of the generated *Pf*CDPK1 model were reasonably accurate. The comparable Ramachandran plot characteristics and RMSD value confirmed the reliability of the 3D-model of *Pf*CDPK1 to be taken further for virtual screening.

**Fig. 1:**
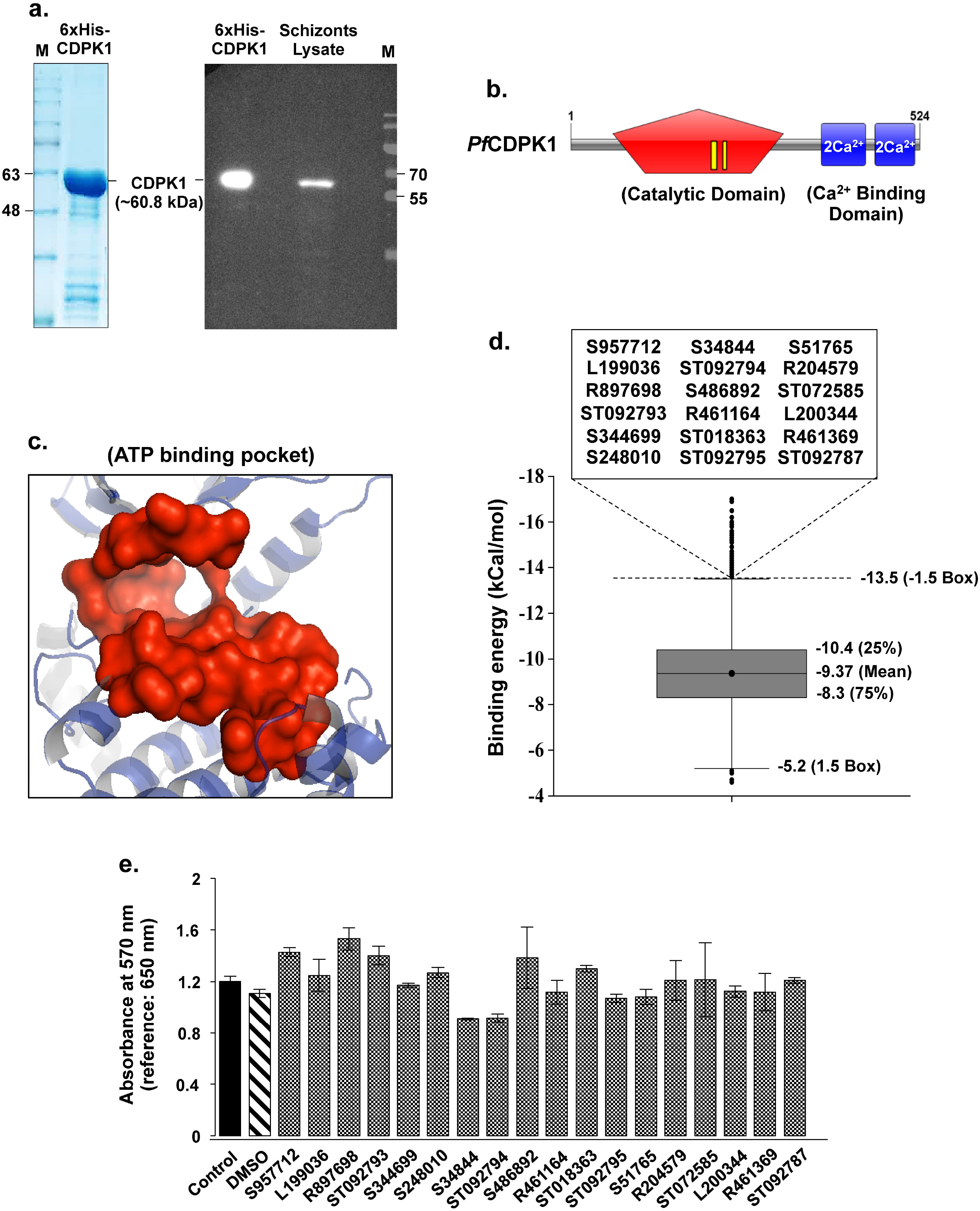
Purified 6xHis-*Pf*CDPK1 protein, three-dimensional structural model of *Pf*CDPK1, virtual screening and cytotoxic potential of compound candidates on HepG2 cell line. **(a)** Coomassie-stained gel showing the level of purity of recombinant *Pf*CDPK1 used for all assays. Anti-sera raised in mice was used to probe recombinantly purified CDPK1 protein and native protein from schizonts lysate. **(b)***Domain architecture of PfCDPK1.* It contains an N-terminal catalytic Kinase Domain (KD, shown in red) tethered to a C-terminal Calmodulin-Like regulatory Domain (CamLD, shown in blue) via auto-regulatory or Junction domain. Activation loop within the KD is shown in yellow. **(c)***Catalytically active ATP binding pocket.* X-Ray diffraction based crystal structure of *Pb*CDPK1 (PDB ID: 3Q5I) was used as template to generate 3D structure of *Pf*CDPK1. **(d)***Virtual screening of MyriaScreen II library against ATP binding pocket of PfCDPK1.* Compounds were ranked on the basis of their negative binding energies along the Y-abscissa. Additional filter was applied to screen for ligands with propensity to follow Lipinski’s rule of five. 18 compounds were finally selected as probable hits and set to submit for experimental validation. **(e)***Cytotoxic potential of the candidate compounds (N=18) obtained from virtual screening of MyriaScreen II library, on HepG2 cell line.* At 100 μM concentration, all 18 compounds displayed negligible cytotoxicity upon 48 hrs of treatment.

### Virtual screening of MyriaScreen II library against ATP binding pocket of CDPK1

MyriaScreen II diversity collection of drug-like screening compounds consists of 10,000 high-purity and diverse molecular candidates from Sigma-Aldrich and TimTec, Inc., created by combining medicinal chemistry and computational expertise, and selected on the basis of diversity and structural relevance for general screening. The collection is largely drug-like according to Lipinski’s rule of five. *Fast docking* of MyriaScreen II library against the generated homology model of CDPK1 was done to computationally screen for ligands with propensity to bind catalytically active ATP binding pocket of the protein (Fig. 1c). Compounds were arranged on the basis of their most negative free binding energies or binding affinity to the ATP binding pocket of the protein, and subsequently ranked by decreasing value of their negative binding energies along the abscissa (Fig. 1d). Compounds were further filtered out on the basis of their propensity to follow Lipinski’s rule of five pertaining to physicochemical properties including Molecular Weight (MW), Octanol/water partition coefficient (LogP), Hydrogen bond acceptors (H-acceptor) and Hydrogen bond donors (H-donors). 18 compounds were finally selected as probable hits and set to submit for experimental validation (Fig. 1d). Besides, all filtered out compounds (N=18) are stabilized by the formation of favourable molecular interactions with amino acid residues constituting ATP binding pocket of CDPK1. Their binding energies, Lipinski’s properties and chemical structures are shown in Table S1.

### Cytotoxic potential of the candidate compounds (N=18) on HepG2 cell line

We further investigated their toxicity on human hepatocellular carcinoma cell line, HepG2 by using MTT cell-viability assay, as mentioned in materials and methods section. At 100 μM concentration, all 18 compounds displayed negligible toxicity on HepG2 cells upon 48 hrs of treatment, with treated cells as healthy as control ones (Fig. 1e).

### Compounds exhibit divergent binding affinities for CDPK1

Compounds (N=18) filtered out from virtual screening of MyriaScreen II library were checked for their interaction with 6xHis-CDPK1 *in vitro* by utilizing SPR. Once immobilized on the SPR sensor chip surface, CDPK1 demonstrated good stability throughout the experiment. With increase in mass concentration of the compounds, gradual increase in SPR sensor signal was observed which linearly correlated with corresponding change in refractive index of the medium immediately adjacent to the SPR sensing surface. The concentration dependent real-time sensorgrams for the interactions, alongwith K_D_ values of each compound are shown in Fig. 2a. The hits displayed a wide range of binding affinities for CDPK1, with K_D_ values varying from 5.4E+04 ± 5.8E+04 μM (ST018363) to 41.2 ± 0.13 μM (ST092794). Notably, interaction with ST092793 produced maximum Response Unit (RU) of around 300.5 at the end of association phase (corresponding to injection of 100 μM concentration of the compound) with K_D_ value of 51.9 ± 0.3 μM. Relative binding affinities of each compound for CDPK1 are depicted in Fig. 2b. Sensor signals obtained from the biomolecular interaction analysis were further transformed into polynomial calibration curves to show relationship between Response at steady state (i.e., at equilibrium, denoted as Req) and concentrations of the compounds (data not shown).

**Fig. 2:**
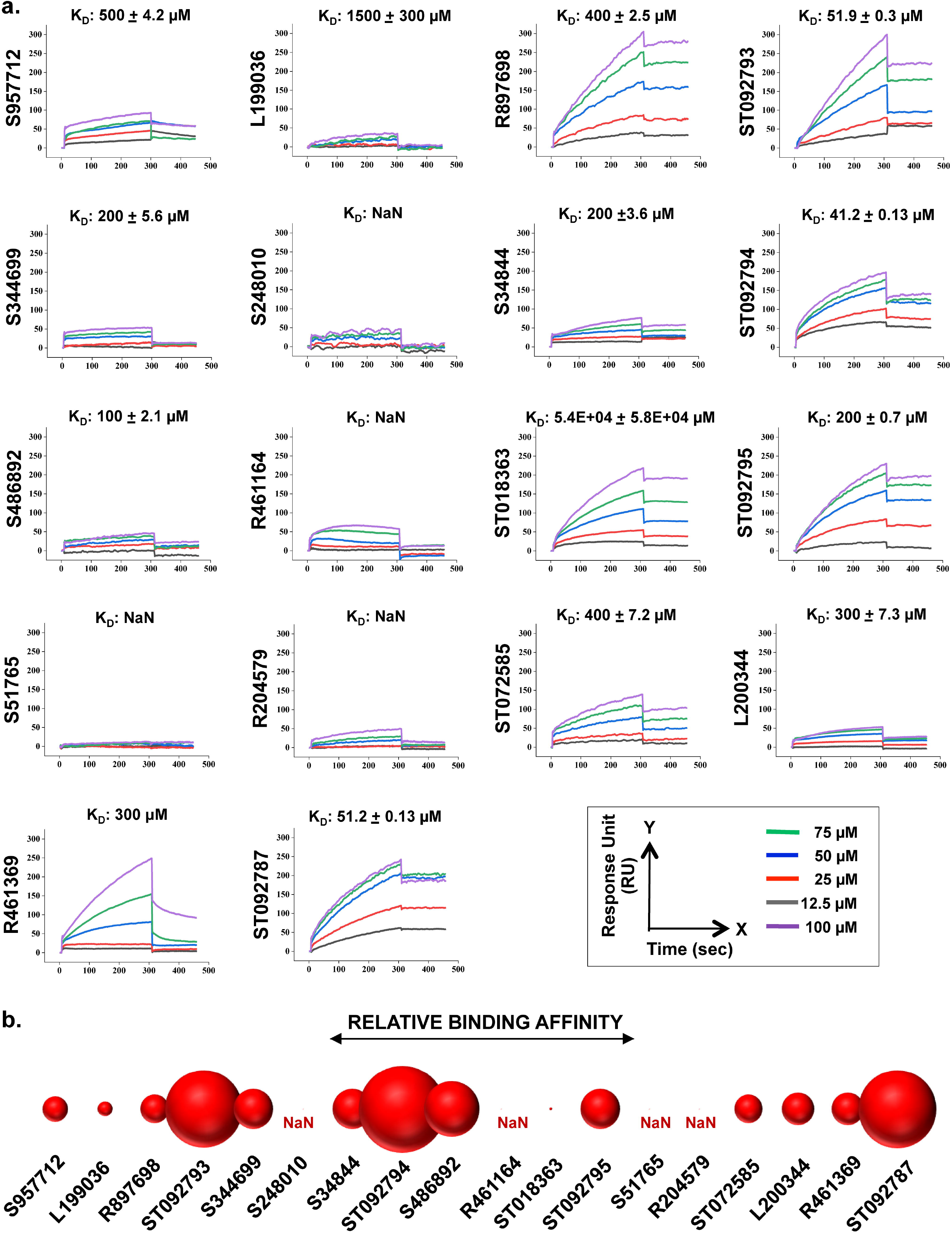
Compounds exhibit divergent binding affinities for CDPK1. **(a)***Concentration dependent real-time sensograms for SPR based biomolecular interaction analysis of CDPK1 and compounds.* Compound candidates procured from virtual screening of MyriaScreen II library displayed a wide range of binding affinities for CDPK1, with K_D_ values ranging from around 5.4 mM (ST018363) to 41.2 μM (ST092794). Interaction with ST092793 (assigned as compound 1) produced maximum RU of approximately 300.5, with K_D_ value of 51.9 μM. **(b)***Relative binding affinities of each compound for CDPK1 are shown*.

SPR analysis suggests that most of the compounds filtered out from virtual screening of MyriaScreen II library are capable of interacting with CDPK1, indicating their probable tendency to inhibit kinase activity of the protein. To verify this inference, the compounds were further examined for their inhibitory effect (if any) on CDPK1 functional activity assays *in vitro* and parasite growth *ex vivo*.

### Compounds 1_(ST092793)_ and 2_(S344699)_ inhibit functional activity of CDPK1 *in vitro*

Since MyriaScreen II library of drug-like small molecular candidates was virtually screened on the basis of probable propensity of the compounds to interact with ATP binding pocket of CDPK1, we utilized *in vitro* kinase assay to demonstrate enzymatic activity of the protein in presence of the filtered out candidates by measuring amount of ADP produced in the kinase reactions. All 18 compounds taken into consideration showed differential inhibition of CDPK1 activity which supports well with our *in silico* virtual screening approach and SPR interaction analysis (Fig. 3a). Notably, two compounds: ST092793 (assigned as compound 1) and S344699 (assigned as compound 2) significantly inhibited phosphorylation activity of CDPK1, accounting for 87.34% ± 0.15% and 68.05% ± 0.48% inhibition, respectively. DMSO, used as solvent for compounds, had no inhibitory effect on the enzymatic reactions and was taken as negative control. Dose-response curves for inhibitory activity of compounds 1_(ST092793)_ and 2_(S344699)_ against CDPK1 protein were found to be sigmoidal, depicting their IC_50_ values to be 33.8 μM and 42.6 μM, respectively (Fig. 3b). Taken together, these results suggest compounds 1_(ST092793)_ & 2_(S344699)_ as potent inhibitors of CDPK1 enzymatic activity.

**Fig. 3:**
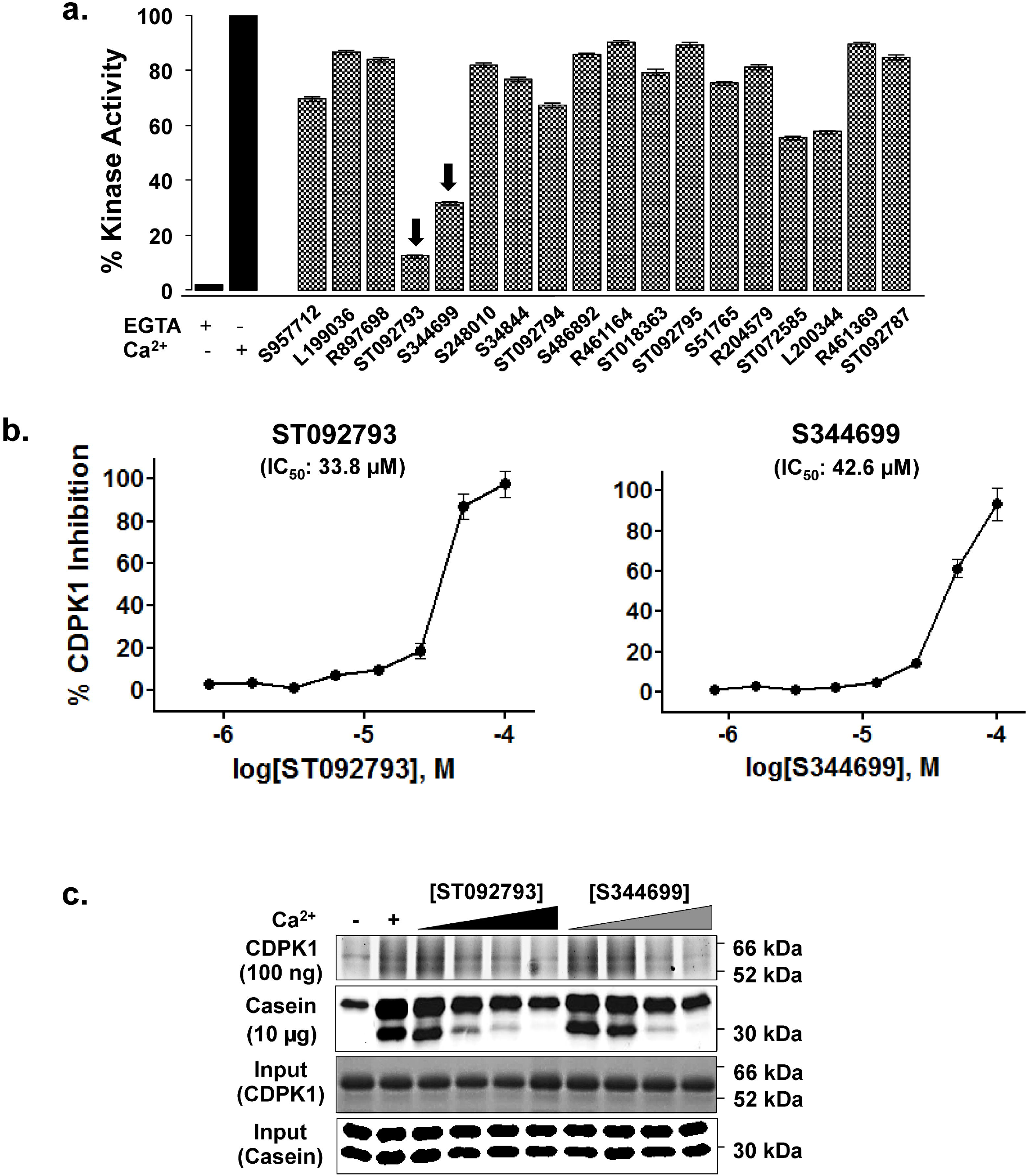
ST092793 (compound 1) and S344699 (compound 2) inhibit functional activity of CDPK1*in vitro*. **(a)** *All 18 compounds showed differential inhibition in CDPK1 activity.* Compounds 1_(ST092793)_ & 2_(S344699)_ significantly inhibited kinase activity, accounting for 87.34% ± 0.15% and 68.05% ± 0.48% inhibition, respectively. DMSO, used as solvent for the compounds, had no inhibitory effect. Enzymatic activity of CDPK1 in control reaction was taken as 100%. **(b)***Dose response curves for inhibitory activity of the lead compounds.* Compounds 1_(ST092793)_ & 2_(S344699)_ exhibited IC_50_ values of 33.8 μM and 42.6 μM, respectively. **(c)***Lead compounds inhibit auto- and trans-phosphorylation status of CDPK1 and dephosphorylated casein, respectively.* Band intensities of CDPK1 (probed with anti-His antibody) and casein in control reactions were taken as 100%. The experiment was done thrice.

Auto-phosphorylation & trans-phosphorylation status of CDPK1 and dephosphorylated casein, respectively, were checked in the absence and presence of varying concentrations of compounds 1_(ST092793)_ and 2_(S344699)_. Fig. 3c shows staining for enzymatic activity, after gel electrophoresis of reaction mixtures, with Pro-Q Diamond phosphoamino acid and Coomassie Brilliant Blue G-250 stains. Decrease in phosphorylation status of both CDPK1 and casein was readily detectable with corresponding increase in concentrations of the compounds, further reaffirming the observation that compounds 1_(ST092793)_ and 2_(S344699)_ are potent inhibitors of CDPK1 activity. We attribute minor differences in apparent size to the gel shift resulting from multiple phosphorylation sites on CDPK1 protein.

### Compounds 1_(ST092793)_ and 2_(S344699)_ inhibit CDPK1 activity by interacting with its ATP binding pocket

Since compounds 1_(ST092793)_ and 2_(S344699)_ inhibit kinase activity of CDPK1 *in vitro*, we wanted to confirm if they accomplish this by directly binding to catalytically active ATP binding pocket of the protein. For this purpose, recombinantly purified CDPK1 was allowed to interact with compound 1_(ST092793)_ and/or 2_(S344699)_, followed by competing this interaction with ATP immobilized on the surface of agarose matrix. Reduction in binding with the ATP-conjugated agarose matrix was observed in the presence of compound 1_(ST092793)_ (91.76%), compound 2_(S344699)_ (86.63%) and in combination of both compounds (89.65%) (Fig. 4a). It was therefore estimated that CDPK1 in complexation with the compound(s) possessed diminished ability to bind ATP, indicating that compounds 1_(ST092793)_ and 2_(S344699)_ inhibit *in vitro* kinase activity of CDPK1 by directly binding to its ATP-binding pocket. Schematic representation of complex formation between catalytically active ATP binding pocket of CDPK1 and compounds 1_(ST092793)_ & 2_(S344699)_ is shown in Fig. 4b.

**Fig. 4:**
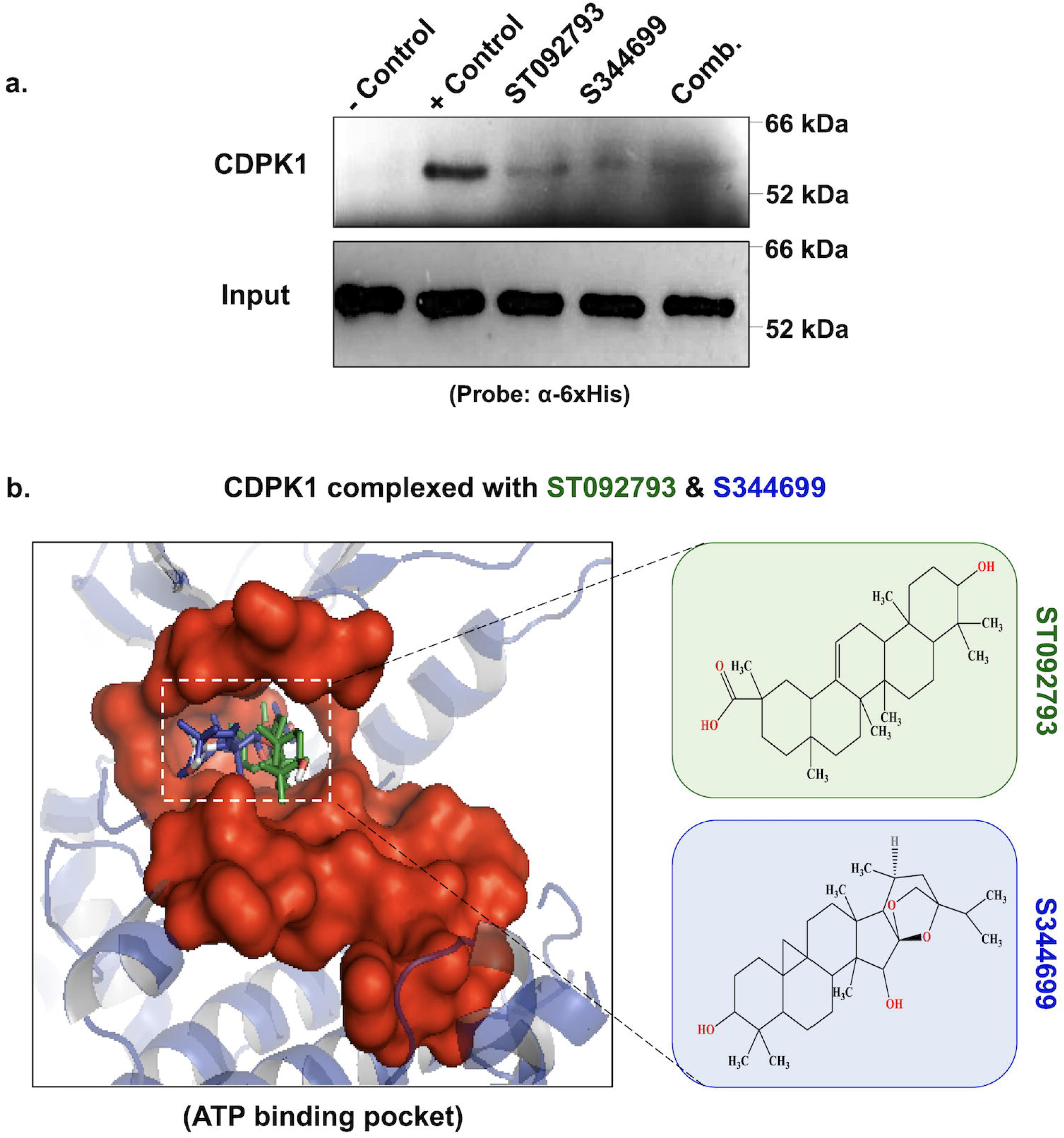
Compounds 1_(ST092793)_ & 2_(S344699)_ inhibit CDPK1 activity by directly interacting with its ATP binding pocket. **(a)***Competitive ATP affinity chromatography assay.* Reduction in binding with the ATP-conjugated agarose matrix was observed upon complexation of CDPK1 with compound 1_(ST092793)_ (91.76%), compound 2_(S344699)_ (86.63%) and in combination of both (89.65%). Band intensity of CDPK1 in untreated reaction was taken as 100%. The experiment was done twice. **(b)***Schematic representation of complex formation between catalytically active ATP binding pocket of CDPK1 and compounds 1_(ST092793)_ & 2_(S344699)_*.

### Compound 1_(ST092793)_ interacts with CDPK1 favourably in the absence of calcium

After affirming CDPK1-inhibitory activity of compounds 1_(ST092793)_ and 2_(S344699)_ *in vitro*, we further sought to establish the knowledge of stoichiometry details and binding modes of complexation between CDPK1 and the *relatively more potent inhibitor*, i.e., compound 1_(ST092793)_, by employing ITC. The representative binding isotherms resulting from titration of compound 1_(ST092793)_ with CDPK1 are represented in Fig. 5a. The compound showed differential binding with CDPK1 in the absence and presence of Ca^2+^. Binding isotherm in the absence of Ca^2+^ was monophasic in nature, reaching a plateau phase at 1:1 stoichiometry (~1.0 molar ratio) which indicates that compound 1_(ST092793)_ interacts with only a single independent binding site (i.e., catalytically active ATP binding pocket) of CDPK1 protein. As further injections continued, gradual decline in exothermic heat resulted in sigmoidal curve ending near zero baseline, indicating saturation of binding sites. On the basis of the nature of the curve, the data were fitted using single-site binding model. In contrast, linear thermogram was observed in the presence of Ca^2+^, indicative of non-specific binding of compound 1_(ST092793)_ with CDPK1. In Fig. 5a, the solid line shows the best fit, and the model reproduces experimental data fairly well. K_a_ values demonstrated that compound 1_(ST092793)_ showed approximately 100 times stronger binding affinity towards CDPK1 in the absence of Ca^2+^ (K_a_ = 3.6 × 10^5^ ± 2.4 × 10^5^M^−1^) than in its presence (K_a_ = 2.6 × 10^3^ ± 25% M^−1^), with the interactions mainly driven by enthalpy: ΔH = −8.1 ± 5.2 kcal·mol^−1^, TΔS = −0.53 kcal·mol^−1^ (± 15-20%) and ΔG = −7.57 kcal·mol^−1^ (± 15-20%). Linear thermogram was obtained in the presence of Ca^2+^, indicative of non-specific and very weak binding of compound 1 with CDPK1.

**Fig. 5:**
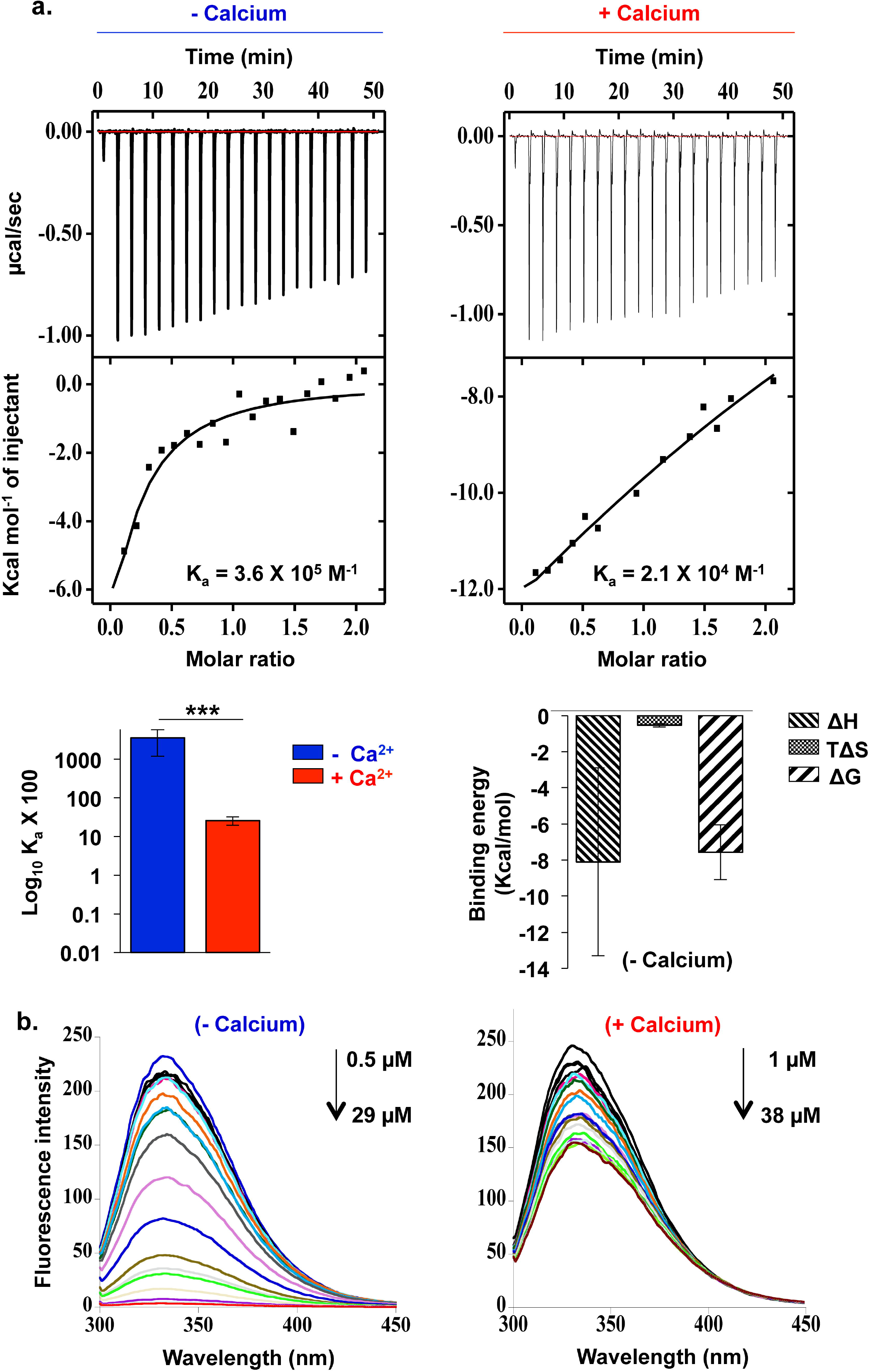
Compound 1_(ST092793)_ interacts with CDPK1 favourably in the absence of calcium. **(a)***Representative ITC binding isotherms resulting from titrations of CDPK1 with compound 1_(ST092793)_.* The compound showed differential binding with CDPK1 in the absence (K_a_ = 3.6 × 10^5^ ± 2.4 × 10^5^ M^−1^) and presence (K_a_ = 2.6 × 10^3^ ± 25% M^−1^) of Ca^2+^. Binding isotherm in the absence of Ca^2+^ reached a plateau phase at 1:1 stoichiometry indicating that compound 1_(ST092793)_ interacts with only a single independent binding site of CDPK1. Also, interaction of compound 1_(ST092793)_ with CDPK1 was enthalpically driven with ΔH = −8.1 + 5.2 kcal·mol^−1^, TΔS = −0.53 kcal·mol^−1^ (± 15-20%) and ΔG = −7.57 (± 15-20%) kcal·mol^−1^. In contrast, linear thermogram was obtained in the presence of Ca^2+^, indicative of non-specific and very weak binding of compound 1 with CDPK1. Solid line represents the best fit of the non-linear experimental data fitted to the single-site binding model. The experiment was done twice. **(b)***Fluorescence spectroscopy supports ITC interaction analysis.* The tryptophan fluorescence maximum for CDPK1 was observed at 333 nm. In the absence of Ca^2+^, titration of CDPK1 with compound 1_(ST092793)_ resulted in sharp fluorescence displacement curves. However, titration in the presence of Ca^2+^ resulted in small variations in the fluorescence emission intensity indicating that compound 1_(ST092793)_ preferentially binds with open or ‘inactive’ conformation of CDPK1. The experiment was done twice.

Overall, the resultant 1:1 stoichiometric complex of compound 1_(ST092793)_ and CDPK1 in the absence of Ca^2+^ was found to be highly stable, supported by stronger K_a_ and favourable binding enthalpy and entropy.

### Fluorescence spectroscopy supports ITC interaction analysis of compound 1_(ST092793)_ and CDPK1

To confirm ITC interaction analysis of compound 1_(ST092793)_ and CDPK1, fluorescence spectroscopic study was performed. Fig. 5b shows typical fluorescence ‘displacement’ titrations, in the absence and presence of Ca^2+^. The tryptophan fluorescence maximum for CDPK1 was observed at 333 nm. In the absence of Ca^2+^, titration of CDPK1 with serial concentrations of compound 1_(ST092793)_ resulted in progressive decline in the tryptophan fluorescence emission intensity throughout the wavelength range of 300–450 nm. In this case, the fluorescence displacement curve was sharp and the compound bound to CDPK1 protein even at lower concentrations. However, in the presence of Ca^2+^, lack of any trend and small variations in the fluorescence emission intensity indicates that compound 1_(ST092793)_ preferentially binds with open or *inactive* conformation of CDPK1, as confirmed by ITC analysis.

### Compound 1_(ST092793)_ confers thermal stability to CDPK1

Thermal Stability Assay (TSA) is a method for detecting target engagement by monitoring thermostability of a given protein in the presence of its ligand. (32) This is based on the principle that target-ligand interactions result in alteration in the thermodynamic parameters of the protein, affecting its stability with corresponding increase in temperature. TSA was optimized for binding analysis between purified recombinant CDPK1 protein and Compound 1_(ST092793)_. Melting curves at temperatures ranging from 40°C to 90°C demonstrated significant thermo-stabilization of CDPK1 in the presence of Ca^2+^ (Fig. 6a). Importantly, in the opted temperature range, CDPK1 treated with Compound 1_(ST092793)_ (100 μM) in the absence of Ca^2+^ showed abundances of the protein higher than the abundances of those treated with DMSO as control, suggesting ligand-dependent thermo-stabilization of CDPK1. No alteration in thermal stability was observed upon treatment of the protein with Compound 1_(ST092793)_ in the presence of Ca^2+^ (Fig. 6b). Our TSA analysis is in consistent with the findings by Scheele, S. *et al* who reported increase in melting Temperature (Tm) of purified CDPK1 from *Toxoplasma gondii*, in the absence of calcium-ions. (33) Further, TSA analysis at 70°C with all 18 compounds (at 100 μM concentration) procured from virtual screening of MyriaScreen II library of small molecules demonstrated that thermal stability of CDPK1 gets relatively more enhanced in the presence of Compound 1_(ST092793)_ (Fig. 6c), further supporting our *in vitro* findings. For quantification of thermostable CDPK1, the signal intensity was normalized to the respective intensity at 40°C.

**Fig. 6:**
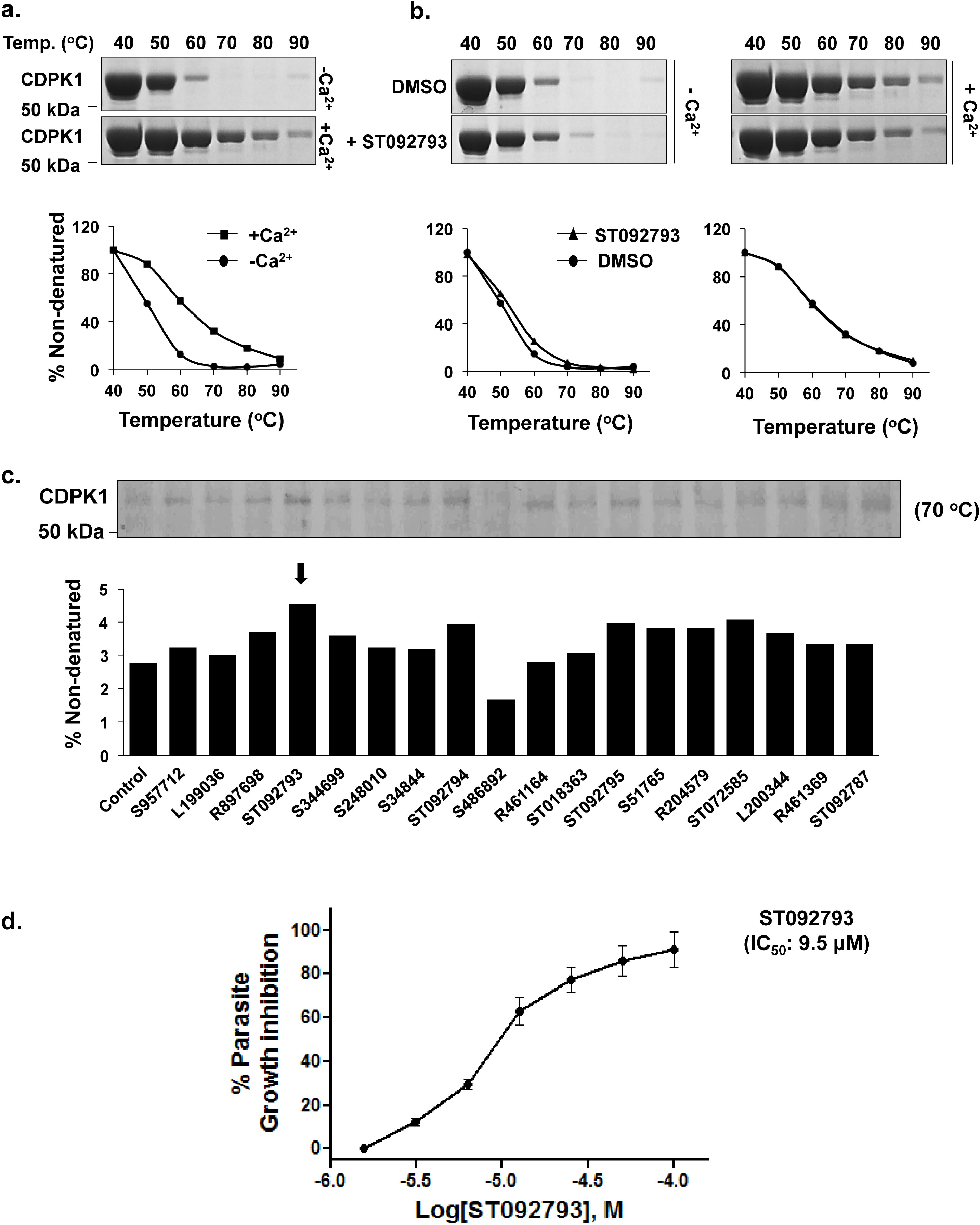
Compound 1_(ST092793)_ confers thermostability to CDPK1. **(a)***Representative melting curves showing thermo-stabilization of CDPK1 in the presence of Ca^2+^.* **(b)***Compound 1_(ST092793)_ mediated thermo-stabilization of CDPK1 is dependent on Ca^2+^ ions.* **(c)***TSA analysis of 18 compounds procured from virtual screening of MyriaScreen II library.* At 70°C, thermal stability of CDPK1 gets relatively most enhanced in the presence of Compound 1_(ST09279)_, further supporting our *in vitro* findings. CDPK1 signal intensity was normalized to the respective intensity at 40°C. **(d)***Evaluation of growth inhibitory effect on the parasite.* Lead compound 1_(ST092793)_ with CDPK1-inhibitory activity *in vitro* showed IC_50_ value of 9.5 μM over 72 hours of intra-erythrocytic life cycle of the parasite.. Parasitemia level of untreated parasites was taken as 100%. X-axis: Log[inhibitor], M; Y-axis: Percent parasite growth.

### Cytotoxic effect of compounds with high selectivity for parasite over host cells

Having observed no cytotoxic effect on HepG2 cell line, lead compound 1_(ST092793)_ with propensity to bind and inhibit the enzymatic activity of CDPK1 was subjected to dose response over 72 hours of intraerythrocytic life cycle of the parasite. Compound 1_(ST092793)_ showed IC_50_ value of 9.5 μM (Fig. 6d). Control parasites were healthy and completed its life cycle producing trophozoites at 72 hrs post-incubation.

## Discussion

In *Plasmodium falciparum*, Ca^2+^-mediated signaling pathways are translated into cellular responses by CDPKs, among which CDPK1 is the most widely studied member of the *cdpk* multigene family (7, 8, 11, 34, 35). Insights into the physiological role of CDPK1 in cellular biology of *P. falciparum* depicts importance of this protein in late schizogony, subsequent invasion of merozoites into host RBCs, formation of male and female gametes, and transmission to the mosquito host (9–12). During gliding and invasion of the parasite, CDPK1 regulates actin-myosin motor by phosphorylating components of IMC, i.e., GAP45 and MTIP. It also regulates cAMP signaling module by phosphorylating and activating PKAr (13). With high abundance, CDPK1 stands alone as a *multistage* signaling regulator of *P falciparum* and depicted as the most sought after molecular target for drug discovery against malaria.

Most of the small molecule *kinase inhibitors* have generally been identified either by high-throughput virtual or fragment-based screening of chemical libraries. Among the drug designing platforms, virtual or molecular *docking* screening utilizes *in silico* robotic platforms for identification of new lead compounds in the process of drug discovery. In this approach, large libraries of unique organic molecules are computationally screened for compounds, collectively referred to as biologically relevant *chemical space*, that complements ‘structural code’ (i.e., ligand binding pocket) of target protein(s), followed by predicting the binding affinity by experimental validation. Filters may be applied to ensure that the hit compounds meet standard of biological relevance or *drug-likeness*, exploting the *druggability* of the target proteins. Once promising candidates are identified, one can utilize medicinal chemistry reactions and synthetic strategies to dynamically generate more diverse compounds around the initial hits, with enhanced efficacy. This whole process of *structure-based* screening and identifying lead candidates frees medicinal chemistry from the tyranny of empirical screening which includes substrate-based design and incremental modification (36–39). Previous studies by, Hou *et al.* identified a phenyl thiadiazole derivative as lead compound by virtual screening against protein kinase CK2, a serine/threonine kinase that has been implicated in pathologies of a variety of human diseases (40). Similarly, Ma *et al.* identified Amentoflavone analogues as Type II inhibitors against JAK2, by structure-based virtual screening of a natural product library, followed by *in silico* optimization and kinetic experiments (41). Further, Su *et al.* identified two potent Rho kinase inhibitors, namely phloretin and baicalein, by integrating high-throughput virtual screening of natural products library with Quantitative Structure-Activity Relationship (QSAR)-based rescoring and kinase assay in order to attenuate Pulmonary Hypertension (PH) (42). Later, Zhao *et al.* identified potential Protein Kinase C (PKC) inhibitors, Fisetin and Tetrahydropapaverine, with activities at nanomolar level, by high-throughput virtual screening against a natural products library to modulate general anaesthetic effects (43).

Earlier attempts have also been made to identify specific inhibitors against *Pf*CDPK1 protein. First evidence came from Kato N. *et al* (2008) who identified a series of structurally related 2,6,9-trisubstituted purines by *in vitro* biochemical screening, by using recombinant *Pf*CDPK1 protein, against a library of 20,000 compounds. Purfalcamine, a representative member of the series resulted in sudden developmental arrest at the late schizont stage in *P. falciparum* (9). Later, Lemercier G. *et al* (2009) identified two distinct chemical series of small molecule inhibitors, compounds **1** and **2** by biochemical screening of 54,000 compounds against recombinant *Pf*CDPK1 protein (44). Also, an earlier study by our group (Bansal A. *et al*, 2013) utilized a peptidic region (P3 peptide; Leu356 to Thr370) of *Pf*CDPK1 Junction Domain (JD) which could specifically inhibit functional activity of the protein, block microneme discharge and erythrocyte invasion by *P. falciparum* merozoites (11). Later, Ansell KH. *et al* (2014) screened a library of 35,000 small molecules by high-throughput *in vitro* biochemical assay followed by chemical optimization to identify chemical series of *Pf*CDPK1-inhibitors (45).

Due to the emergence of parasite resistance to existing frontline antimalarial drugs, there is a pressing need to identify and characterize *new* anti-malarial molecules that would target the parasite at multiple proliferative stages during its life cycle. In this context, since *Pf*CDPK1 has been previously identified as one such target, we have utilized a structurally diverse collection of 10,000 drug-like molecules from MyriaScreen II library to screen for ligands complementing ATP binding pocket of CDPK1, as represented schematically in Fig. 7. In the absence of an experimentally determined atomic structure of *Pf*CDPK1, earlier attempts have been made to resolve homology modeling based three-dimensional structure of *Pf*CDPK1 (9, 11, 44). In all these studies, *distantly* related orthologous sequences of *Pf*CDPK1 either from humans (death-associated protein kinase, PDB ID: 1JKK) (9) or from another apicomplexan parasite, *Cryptosporidium parvum* (calcium/calmodulin-dependent protein kinase, PDB ID: 2qg5; and, *Cp*CDPK3, PDB ID: 3LIJ) (11, 44) was used as template for structural modeling of *Pf*CDPK1. Owing to the high sequence identity (~93%) between CDPK1 orthologs from *P. falciparum* strain 3D7 and evolutionarily *closely* related species, *P. berghei* strain ANKA, we used X-Ray diffraction based structural model of *Pb*CDPK1 (the most recently resolved crystal structure of CDPK1 protein, PDB ID: 3Q5I) as template to generate 3D structural coordinates of *Pf*CDPK1. Stereochemical quality and accuracy of the structurally refined model of *Pf*CDPK1 (Fig. S1a and S1b) indicated that the backbone dihedral angles: phi and psi were reasonably accurate, thus confirming reliability of the generated model for performing docking-based virtual screening (Fig. S1c). The candidates were filtered and ranked from MyriaScreen II library on the basis of their most negative free binding energies or binding affinity towards ATP binding pocket of the protein (Fig. 1c). After applying additional filter of Lipinski’s rule of five, 18 compounds were finally selected as probable hits and set to submit for experimental validation (Fig. 1d and Table S1). Also, at 100 μM concentration, all 18 compounds displayed negligible toxicity on HepG2 cells upon 48 hrs of treatment, with treated cells similar to untreated healthy control (Fig. 1e). Real-time biomolecular interaction analysis was carried out with SPR to determine and compare respective binding strengths of the hits (N=18) and recombinantly purified 6x-His-CDPK1 protein, at physiologically relevant concentrations. Wide range of binding affinities was observed, with K_D_ values ranging from around 54 mM (ST018363) to 41.2 μM (ST092794). Compound 1 _(ST092793)_ showed K_D_ value of 51.9 ± 0.3 μM, generating maximum RU of around 300.5 (Fig. 2).

**Fig. 7:**
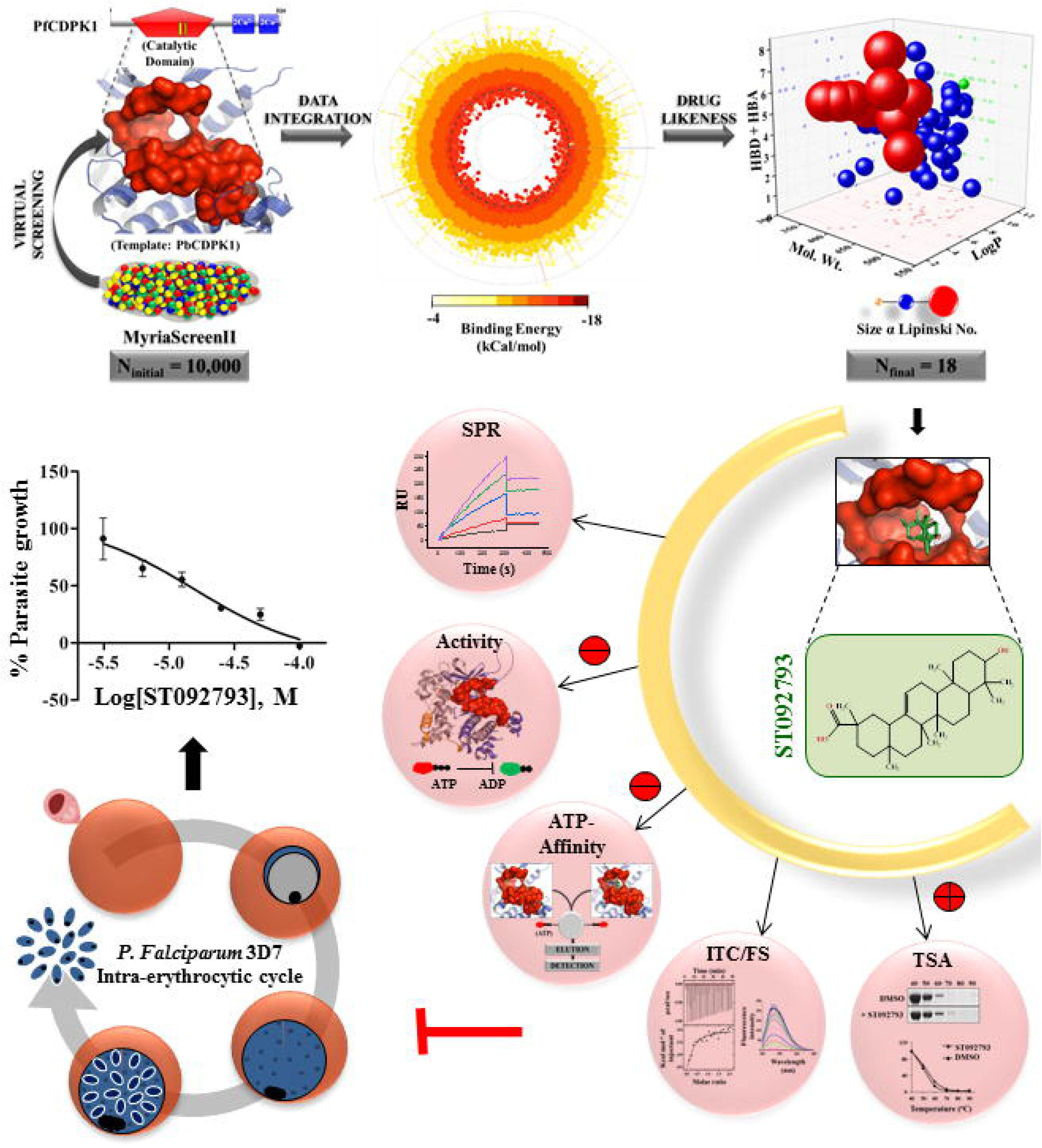
Overall schematic representation. Structurally diverse collection of 10,000 drug-like molecules from MyriaScreen II library was utilized to screen for ligands complementing ATP binding pocket of CDPK1, thus inhibiting parasite growth *in vitro*.

To verify probable tendency of these compounds to inhibit functional activity of the kinase, compounds (N=18) were further subjected to CDPK1 functional activity assays *in vitro*. All 18 compounds showed differential inhibition of CDPK1 activity at 50 μM concentrations, with compounds 1_(ST092793)_ and 2_(S344699)_ accounting for 87.34% ± 0.15% and 68.05% ± 0.48% inhibition, respectively (Fig. 3a). Dose-response curves for CDPK1-inhibitory activity of compound 1_(ST092793)_ or 2_(S344699)_ were found to be sigmoidal, depicting their IC_50_ values to be 33.8 μM and 42.6 μM, respectively (Fig. 3b). Also, decrease in phosphorylation status of CDPK1 and casein was observed with increase in concentrations of both of these compounds (Fig. 3c), which suggests that compounds 1_(ST092793)_ and 2_(S344699)_ are potent inhibitors of CDPK1 protein. To further authenticate, if compounds 1_(ST092793)_ and 2_(S344699)_ inhibit functional activity of CDPK1 by directly competing with ATP for binding to catalytically active pocket of the protein, affinity chromatography of CDPK1 pre-incubated with compound 1_(ST092793)_ and/or 2_(S344699)_ was performed. By doing so, reduction in binding to ATP-conjugated agarose matrix was observed in the presence of compounds 1_(ST092793)_ (91.76%) and 2_(S344699)_ (86.63%), indicating that both of these compounds directly bind to catalytically active ATP-binding pocket of CDPK1 and thus inhibits its enzymatic activity (Fig. 4).

Binding affinity and stoichiometry details of the complex formation between *relatively more potent* compound 1_(ST092793)_ and CDPK1 was further examined by utilizing ITC, in the absence and presence of Ca^2+^. K_a_ values demonstrated that compound 1_(ST092793)_ showed higher binding affinity towards CDPK1 in the absence of Ca^2+^ (K_a_ = 3.6 × 10^5^ M^−1^) than in its presence (K_a_ = 2.1 × 10^4^ M^−1^), with the interactions mainly driven by enthalpy (ΔH = −8.1 kcal.mol^−1^). Binding isotherm in the absence of Ca^2+^ reached a plateau phase at 1:1 molar ratio, indicating that the compound interacts with only a single independent binding site (ATP binding pocket) of the protein. In contrast, in the presence of Ca^2+^, non-specific interaction was observed as depicted by the linear isotherm (Fig. 5a). ITC interaction analysis was further confirmed by measuring intrinsic tryptophan fluorescence through spectroscopic studies in the presence of increasing concentrations of compound 1_(ST092793)_. Fluorescence displacement titrations in the absence of Ca^2+^ resulted in progressive decline in the tryptophan fluorescence intensity. However, lack of any trend in the presence of Ca^2+^ indicated that compound 1_(ST092793)_ preferentially binds with CDPK1 in its open or ‘inactive’ conformation, as confirmed by ITC (Fig. 5b).

To further confirm the target engagement, TSA was applied to monitor thermostability of CDPK1 upon treatment with compound 1 _(ST092793)_, in the absence and presence of Ca^2+^. Melting curves at temperatures ranging from 40°C to 90°C demonstrated thermo-stabilization of CDPK1 in the presence of Ca^2+^ (Fig. 6a). More importantly, treatment of CDPK1 with Compound 1 _(ST092793)_ resulted in enhanced thermo-stabilization of the protein in a Ca^2+^ dependent manner (Fig. 6b). Further, TSA analysis with all 18 compounds procured from virtual screening of MyriaScreen II library demonstrated that thermal stability of CDPK1 gets relatively highly enhanced in the presence of Compound 1 _(ST092793)_, further supporting our *in vitro* findings (Fig. 6c). Finally, lead compound 1_(ST092793)_ obtained from virtual screening of MyriaScreen II library was subjected to dose response over 72 hours of intra-erythrocytic life cycle of the parasite. Compounds 1_(ST092793)_ with CDPK1 inhibitory activity *in vitro* showed IC_50_ value of 9.5μM (Fig. 6d).

In conclusion, we adopted a structure-based virtual chemical library screening approach in combination with extensive biochemical and biophysical characterization based tools to identify novel lead candidates, Compound 1_(ST092793)_ and Compound 2_(S344699)_ capable of inhibiting CDPK1 activity *in vitro*. The inhibitory potency of these compounds was authenticated by favorable interactions with the catalytically active ATP binding pocket of the protein. These compounds exhibited potent cytotoxicity against malaria parasite *ex vivo*, indicating their promising candidacy for malaria treatment. Further studies can be planned to improve CDPK1 inhibitory activity and kinase selectivity of these compounds by designing new analogues of the same. Altogether, our study proposes a novel CDPK1-selective inhibitor with strong anti-malarial activity and represents a combinatorial high-throughput platform involving *in silico* and *in vitro* applications as a blueprint for the rational identification of novel selective inhibitors.

## Abbreviations

CDPK1: Ca^2+^-Dependent Protein Kinase 1
ACT: Artemisinin-based Combinatorial Therapy
MDR: Multi-Drug Resistance
GMS: Greater Mekong Sub-region
CDPKs: Calcium-Dependent Protein Kinases
KD: Kinase Domain
CamLD: Calmodulin-Like regulatory Domain
J: Junction domain
CaMKs: Calmodulin-dependent protein Kinases
PKG: Protein Kinase G
IMC: Inner Membrane Complex
GAP-45: 50-kDa Glideosome-Associated Protein
MTIP: Myosin Tail domain Interacting Protein
PKAr: regulatory subunit of cAMP-dependent Protein Kinase A
AMA-1: Apical Membrane Antigen-1
EBA-175: 175 kDa Erythrocyte Binding Antigen
RMSD: Root-Mean-Square Deviation
MW: Molecular Weight
LogP: Octanol/water partition coefficient
H-acceptor: Hydrogen bond acceptors
H-donors: Hydrogen bond donors
RU: Response Unit
Ka: Binding affinity constant
ΔH: enthalpy change
N: stoichiometry
ΔG: Gibbs free energy change
ΔS: entropy change
TSA: Thermal Stability Assay
BI: Boehringer Ingelheim
PGVL: Pfizer Global Virtual Library
AZ-VL: AZ-Virtual Library
QSAR: Quantitative Structure-Activity Relationship
PH: Pulmonary Hypertension
PKC: Protein Kinase C
AML: Acute Myeloid Leukemia
FLT3: FMS-like Tyrosine Kinase 3
PDB: Protein Data Bank
SDF: Structural Data Format
CDD: Conserved Domain Database
DruLiTo: Drug Likeness Tool
CDK: Chemistry Development Kit
IPTG: IsoPropyl ß-D-1-ThioGalactopyranoside
OD: Optical Density
EDTA: Ethylene Diamine Tetraacetic Acid
ßME: ß-MercaptoEthanol
PIC: Protease Inhibitor Cocktail
SPR: Surface Plasmon Resonance
Ka: Association rate constant
Kd: Dissociation rate constant
RT: room temperature
NHS: N-HydroxySuccinimide
NAS: N-HydroxySuccinimide esters
IRL: Integrated Rate Law
kd: intercept
ka: slope
DTT: DiThioThreitol
8AH-ATP-agarose: {8-[(6-Amino) Hexyl]-amino-ATP-agarose
ITC: Isothermal Titration Calorimetry
FS: Fluorescence Spectroscopy
TSA: Thermal Stability Assay
CETSA: Cellular Thermal Shift Assay
EtBr: Ethidium Bromide
FL-2: Fluorescence signal
MTT: [(3-[4,5-dimethylthiazol-2-yl]-2,5-diphenyl tetrazolium bromide)]
SD: standard deviation
Req: Response at equilibrium

## Acknowledgements

This work has been funded by DST-EMR from the Department of Science and Technology (DST), Ministry of Science and Technology, Government of India. Shailja Singh is a recipient of the Innovative Young Biotechnologist Award (IYBA) and National Bioscientist Award from Department of Biotechnology (DBT). RJ is supported from University Grants Commission-Junior Research Fellowship (UGC-JRF). Authors acknowledge SPR facility of Advanced Instrumentation Research Facility (AIRF), Jawaharlal Nehru University (JNU), and central facility of Special Center for Molecular Medicine (SCMM), JNU for flow cytometry. The lab facilities of Shiv Nadar University and SCMM, JNU are also acknowledged.

## Author Contributions

RJ and SS designed the experiments. RJ and SG performed the experiments. SS contributed reagents. RJ, MM, SS analyzed data. RJ and SS wrote the manuscript.

## Notes

The authors declare no competing financial interest.

