## Supplementary file for "Development of novel anti-malarial from structurally diverse library of molecules, targeting plant-like Calcium Dependent Protein Kinase 1, a multistage growth regulator of *P. falciparum*"

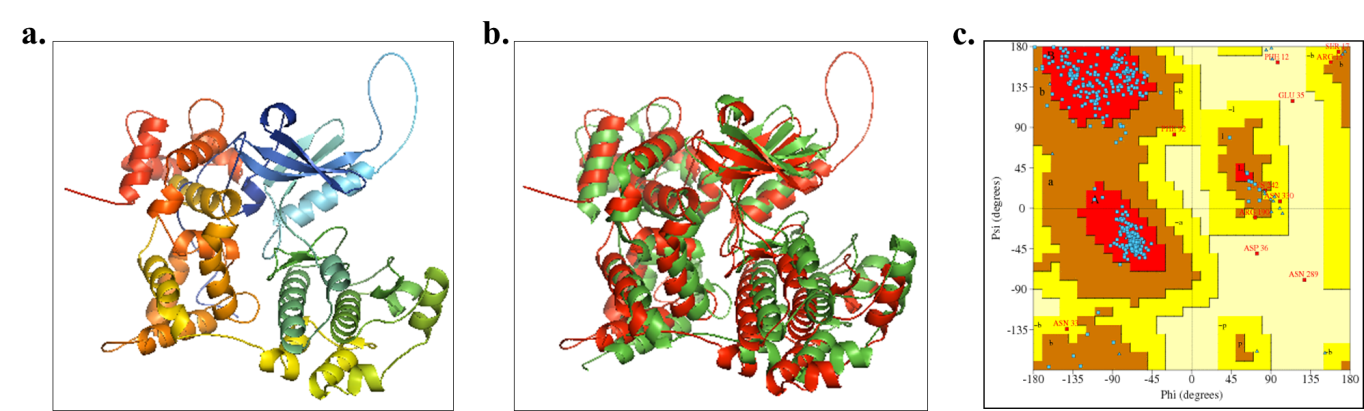

**Fig. S1: Homology model of *Pf*CDPK1 and its quality assessment. (a)** *Comparative structural model of PfCDPK1.* Based on high sequence similarity (93%) between *Pf*CDPK1 and *Pb*CDPK1, X-Ray diffraction based structural model for *Pb*CDPK1 (PDB ID: 3Q5I) was used as template to model 3D structure of PfCDPK1. **(b)** *Optimal rigid-body superimposition of PbCDPK1 and PfCDPK1.* Overall RMSD value of the C-alpha atomic coordinates was found to be 0.31 Å, suggesting a reliable modeled structure of *Pf*CDPK1. **(c)** *Stereochemical assessment of the generated homology model.* Assessment of backbone dihedral (torsion) angles: phi (Ø) and psi (Ψ) of the amino acid residues displayed 89.2% of the residues lying in the most favored (“core”) regions, with 8.1%, 1.9%, and 0.8% residues in “additional allowed”, “generously allowed” and “disallowed regions” of Ramachandran plot, respectively (Figure 1).

**Table S1: Binding energies, Lipinski’s properties and chemical structures of the compound candidates (N = 18) procured from MyriaScreen II library. (Provided as separate excel sheet, labeled as Table S1)**
